# Untargeted and Semi-Targeted Metabolomics Approach for Profiling Small Intestinal and Fecal Metabolome Using High-Resolution Mass Spectrometry

**DOI:** 10.1101/2024.09.16.613180

**Authors:** Alexandre Tronel, Morgane Roger-Margueritat, Caroline Plazy, Valérie Cunin, Ipsita Mohanty, Pieter C. Dorrestein, Thomas Soranzo, Audrey Le Gouellec

## Abstract

The gut microbiome is a complex ecosystem varying along different gut sections, consisting of metabolites from food, host, and microbes. Microbially-derived metabolites like bile acids and short-chain fatty acids interact with host physiology. Current studies often use fecal samples, which don’t fully represent the upper gut due to stratification. To sample the proximal gut microbiome, endoscopic methods or new non-invasive devices are used. We developed an approach combining untargeted and semi-targeted metabolomics using a Q-Exactive Plus Orbitrap mass spectrometer to profile gut metabolites. We initially selected 49 key metabolites based on specific criteria, validated them through repeatability tests, and created a compound database with TraceFinder software. Our workflow enables molecule annotation in untargeted studies while validating 37 metabolites in semitargeted analyses. This method, applied to clinical trial samples (NCT05477069), shows promise in discovering new gut metabolites.

**Highlights:**

- **Innovative Combined Metabolomics Workflow:** The study introduces a combined approach that integrates semi-targeted and untargeted metabolomics analyses to characterized small intestinal and fecal metabolomes. This method allows for the relative quantification of carefully selected metabolites and the reanalysis of these metabolites using evolving curated databases, enhancing the understanding of the gut microbiome-health axis.
- **Untargeted Metabolomics Strategy:** The untargeted approach aims to determine the global metabolic profile of samples and discover new metabolites. This involves processing data through a detailed pipeline, statistical analysis, and feature annotation using tools like MZmine, MetaboAnalyst, and the GNPS platform.
- **Proof of Concept on Clinical Samples:** The combined approach was tested on clinical samples from a participant in a clinical trial, revealing distinct metabolomes between small intestinal content and fecal samples. This proof of concept demonstrated the method’s ability to identify and quantify metabolites, showing significant differences in metabolite abundance between the two sample types and highlighting the potential for discovering new bile acids through molecular networking.

## 1. Introduction

The gut microbiome, including bacteria, fungi, viruses, protists, and archaea, along with their biological components (metabolites, proteins, and free nucleic acids [1]), is modulated by environmental factors including diet [2], medication [3] and host physiology. And the gastrointestinal tract (GIT) is characterized by distinct environmental conditions that play a crucial role in shaping the microbiome along its length. Various gradients, such as pH, oxygen and mucus [4], directly influence the microbiome composition [5] from the beginning to the end of the GIT, resulting in an anoxic environment in the distal gut. Additionally, host secretions significantly impact microbial composition. These secretions, including gastric juices, pancreatic fluids, and bile, contain enzymes and metabolites that exert selective pressure on microbes. Bile acids (BAs), derived from the host, primarily function to emulsify lipids, enhancing their solubility and absorption in the small intestine [6]. However, BAs can also cause bacterial membrane and DNA damages, and oxidative stress due to their detergent properties [7]. Bacteria have developed mechanisms to modify BAs to survive in the environment [8]. Furthermore, the host immune system shapes the microbiome by secreting antimicrobial peptides and secretory immunoglobulin A into the lumen. Beyond the spatial environmental impacts, temporal factors, particularly transit time, significantly influence the composition of the microbiome [9]. Consequently, the human gut microbiome exhibits a stratified distribution from the duodenum to the colon, characterized by a progressive increase in bacterial density and diversity along the gut [10]. While the fecal microbiome has been extensively studied, it does not represent the microbiome of the proximal intestine. The small intestinal microbiome remains less characterized due to limited accessibility, as many collection methods are invasive and require hospitalization, such as surgery [11]. However, advancements in medical device (MD) development for non-invasive collection of intestinal liquid samples offer significant opportunities to study the small intestinal microbiome [12] as recently demonstrated in several clinical investigations [13–16].

So far, only a few studies, such as the one by Shalon *et al*., have investigated the small intestinal metabolome by using MD to collect intestinal liquid. The authors conducted untargeted metabolomics on small intestinal liquid samples and fecal samples [13,17]. They identified 22 microbially conjugated BAs in the small intestinal liquid samples of all participants, with significantly higher concentrations compared to fecal samples [13]. Targeted metabolomics approaches on BAs and short-chain fatty acids (SCFA) revealed notable differences between feces and small intestinal samples, with SCFAs exhibiting increased levels in the distal parts of the gut [17]. Other studies employing various sampling methods have investigated the small intestinal metabolome [16,18–20]. Studying the small intestine metabolome is of great interest, particularly because the proximal gut is where nutrient absorption occurs. As previously mentioned, numerous host secretions are released in the duodenum to facilitate food degradation. The gut microbiome plays several crucial roles and is associated with four major functions: metabolic, immune, endocrine, and regulation of the gut-brain axis. Most interactions between microbes, and between the host and microbes, are mediated by metabolites. Gut microbiota-derived metabolites can originate directly from the diet, from the host (such as amino–acids-conjugated bile acids), or be synthesized de novo [21]. It has been demonstrated, for instance, that SCFAs can directly modulate gut permeability [22] and the immune system [23]. Consequently, it is crucial to better understand and characterize the metabolome of the GIT. Metabolomics, one of the most recent omics fields, relies on advanced high-throughput analytical chemistry techniques such as mass spectrometry coupled with liquid chromatography (LC-MS) or nuclear magnetic resonance (NMR) spectroscopy [24]. The advancements in analytical techniques and bioinformatics have enabled the study of complex samples and the analysis of vast amounts of data. There are three main approaches to study the metabolome [25]. The first approach, targeted metabolomics, involves measuring specific metabolites (usually tens not hundreds), allowing for the absolute quantification of these molecules [26]. The second approach, untargeted metabolomics, employs an unbiased method for global metabolic footprint determination and discovery [27]. Untargeted metabolomics are exploratory in nature, detecting a wide array of metabolites without prior selection and reporting their relative amounts based on normalized peak areas. The third approach is a combination of the first two: semi-targeted metabolomics. In this approach, data acquisition follows an untargeted method, but pure compounds are pre-analyzed to ensure the detection and determine compound characteristics. This allows analyzing numerous metabolites with some predefined targets and reporting a combination of normalized peak areas and absolute concentrations [26]. Each assay type serves distinct purposes and has specific validation requirements, thereby contributing uniquely to the field of metabolomics research.

To gain insight into the biologically significant properties of the human GIT microbiome, it is essential to have validated analytical measurements that precisely describe various characteristics of the microbial community, both quantitatively and qualitatively. In this article, we describe the process we employed to develop a combined untargeted and semi-targeted metabolomics approach to analyze small intestinal and fecal samples using LC-MS/MS. Initially, we identified 49 metabolites of interest in both matrices, including BAs, amino acids, and vitamins, before proceeding with the development of the semi-targeted approach. For semi-targeted metabolomics, we created a compound database using TraceFinder4 software and used it to detect the metabolites of interest. To ensure accurate detection, we evaluated the coefficient of variation and the linearity of each metabolite. Our approach enabled us to detect and quantify 37 metabolites using our semi-targeted method, while also determining the global metabolite profile and discovering new molecules with the untargeted method.

## 2. Materials and Methods

### Metabolomic workflow construction

This metabolomic workflow was built to answer different biological questions. First, we wanted to determine the small intestinal and fecal metabolomes using untargeted metabolomics. Then, using this unbiased approach one of the objectives was to identify special metabolites from each matrix. To better identify and to semi-quantify the metabolites, we added a semi-targeted approach based on 49 selected compounds susceptible to discriminate samples from their origin.

### 2.1. Authentic chemical standard collection

The authentic chemical standard collection is composed of individually and commercially pure compounds (e.g. Merck). Each standard was weighted and resuspended for solid metabolites or diluted for liquid metabolites, in pure ULC/MS-CC/SFC water (Biosolve Chimie) or ULC/MS-CC/SFC absolute methanol (Biosolve Chimie) at the final concentration 10mM or 1mM depending on the solubility limit. The solubilized standards were banked and stored at −80°C. For the mix of metabolites, each metabolite was added to achieve a final concentration of 100µM after a nitrogen evaporation cycle. Deuterated standards (L-Phenylalanine-3,3-d2 / L-Leucine-3,3-d2 / DL-Tryptophan-2,3,3-d3) were purchased from CLUZEAU INFO LABO (FRANCE).

### 2.2. LC-MS/MS method

#### Instrumental and chromatographic settings

As described by Roca *et al*., most of the fecal metabolites are polar or semi-polar compounds [28]. A LC-MS/MS method using a polar C18 column (Reverse Phase) was performed to capture a maximum of the metabolomic footprint.

Untargeted and semi-targeted metabolomics approaches were conducted using an Ultra-high-performance liquid chromatography (Vanquish Flex, ThermoFisher Scientific, Waltham, MA, USA) coupled with a Q Exactive Plus Orbitrap high resolution mass spectrometer (ThermoFisher Scientific, Waltham, MA, USA).

The chromatographic separation was carried out on a Luna Omega polar C18 (2.1mm × 150 mm, 1.6 μm, Phenomenex, Torrance, CA, USA) at 30 °C with a flow elution rate of 400 μL/min. The temperature of the autosampler compartment was set at 4°C, and the injection volume was 5μL. For the chromatographic settings, the mobile phases consisted of A (HPLC water + 0.1% formic acid) and B (HPLC acetonitrile (ACN) + 0.1% formic acid). Elution started with an isocratic step of 2 min at 1% mobile phase B, followed by a linear gradient from 1% to 100% mobile phase B for the next 12 min. The next 6 min were with 100% mobile phase B before returning to 1% B for 5 min corresponding to the column equilibration. The mass spectrometer was fitted with an electrospray source (ESI) operating in positive and negative ionization modes. The scan range was from *m/z* 85.0 to 1275.0 with a resolution of 70,000 at a *m/z* 200. MS2 fragmentation was performed at three collision energies (CEs: 10, 35, 55 eVs) on the top 10 of the most abundant parent ions during full scan. Quality controls (QC) were used to ensure the quality of the LC-MS/MS acquisition. Deuterated compounds were used for controlling extraction process (d-leucine) and injection (d-tryptophan and d-phenylalanine). Pooled QC assembled from all biological samples ensured the stability of peak detection and intensity. They were injected every 10 injections. Diluted pooled QC (1/2, 1/4, 1/8, 1/16) were used to evaluate the linearity of detection of features.

#### Sample preparation for metabolomics

-Two distinct matrices were prepared for LC-MS/MS acquisition: intestinal liquid and feces. Samples were then homogenized (vortex and sonication 10 min). MeOH spike with d-leucine were added in a ratio of 1:4 (w/v) for metabolites extraction and protein precipitation. The next steps of the protocol are the same as the intestinal liquid preparation. For intestinal liquid, metabolites were extracted by addition of cold methanol (1:4 (v/v)) spiked with d-leucine for protein precipitation and vortexed for 20s. Samples were incubated on ice for 30 min and centrifuged 15 min, 15,000g, +4°C. Supernatants were separated from pellets and evaporated (nitrogen evaporation). Dry extracts were resuspended with UHLPC solvent (80% water, 20% MeOH, 1% ACN, 0.1% formic acid and d-Trp/d-Phe at 5µM).

### 2.3. m/z-RT and MS/MS reference library

To match each extracted-ion chromatogram (EIC) peaks, we used the Xcalibur software (ThermoFisher Scientific). For each metabolite, we assigned a retention time. MS/MS fragmentation for each metabolite was determined using the Xcalibur based on parent ions. Experimental fragments were compared to MassBank spectra for validation.

### 2.4. Compound database construction on TraceFinder

The compound database was built using TraceFinder 4.1 General Quan (ThermoFisher Scientific). In this database, metabolite names, chemical formula, *m/z* ratio, RT, ionization mode and adducts were registered. We also added fragments obtained from MS2 to validate the molecules. To build the master method, a raw file containing all the metabolites of interest was implemented. The detection algorithm used was Genesis.

### 2.5. LC-MS/MS experimental validation

#### Repeatability

- Repeatability was evaluated by calculating the coefficient of variation (CV) for each metabolite of interest. In order to do so, pooled metabolites were injected 10 times and peak intensities were determined using TraceFinder4. The CV was calculated from the standard deviation of the 10 peak intensities divided by the mean intensity of each metabolite, multiplied by 100.

#### Linearity of the detection

- To evaluate the linearity of detection for our 41 metabolites, we tested a concentration range from 1µM to 100µM on pooled metabolites in triplicate. The peak areas were extracted using TraceFinder4 and the output was treated in a table with Python v. 3.9.5, precisely Pandas module (v. 1.4.0). The areas were then plotted against the corresponding concentration for each metabolite thanks to Matplotlib v. 3.5.1. Finally, a linear regression and the associated R-squared (R^2^) were obtained with the Scipy v. 1.13.1 module from Python.

### 2.6. Data analysis

Untargeted metabolomics data were analyzed using MzMine3 [29]. Briefly after the conversion of raw files in .mzXML files using MSconvert, data were loaded on MZmine3 [29] following the data processing procedure [30]. The noise threshold on our instrument was evaluated in full scan at an intensity of 1E6 and 1E3 in MS2. Chromatograms were built with ADAP chromatogram builder [31] and deconvolution was performed with the local minimum feature resolver. After isotope grouping, the aligned feature table was used for further data curation. Briefly, using Excel, features found in both blank and samples were deleted. Features with a CV > 30% in pooled QC and with a linearity < 0.7 in diluted pooled samples were also deleted from analysis. For features annotation, we used GNPS (Global Natural Products Social Networking) [32], Sirius 6.0.1 [33] and performed molecular networking with Cytoscape3.10.2.

For semi-targeted metabolomics, raw data were directly analyzed using TraceFinder4 and the previously compound database. Different levels of confidence were given to the metabolites of interest depending on the validation with MS2 spectrum [34]. Indeed, only the first 10 features with the highest intensity were fragmented to determine the MS2 spectrum. If the MS2 spectrum matched with the fragments included in the compound database, the level of consistency was assigned as level 1. If there is no scan, the level assigned was 2. Once identifications were validated, peak areas were extracted and analyzed

After the standard linearity, the analysis performed on intestinal liquid and feces was treated with TraceFinder4 to obtain peak areas. To assess sample losses due to the metabolite extraction method, the recovery percentage was measured by dividing the detected d-leucine in each sample by the expected d-leucine quantity. Then the following formula was applied to obtain metabolite quantity in arbitrary unit per mg for feces or µL for intestinal liquid: peak area / (recovery*mg for feces or µL injected). Finally, for each metabolite, the ratio between SI and feces content could be computed to evaluate in which matrix the metabolite of interest was the most abundant. If the metabolite was totally absent in one of the matrices, the ratio could not be calculated.

BA annotations were achieved using Feature Based Molecular Networking (FBMN) on GNPS2 and an expanded set of bile acid libraries [32,35,36]. The GNPS2 job can be accessed using <u>the link</u>.

## 3. Results

### 3.1. Design of the combined Metabolomics semi-targeted and untargeted analytic Workflow

In recent years, fecal and small intestinal liquid samples have become the preferred matrices for studying the gut microbiome-health axis thanks to the development of non-invasive collection methods and their unique reflection of an individual’s lifestyle. The continuous advancements in science and our understanding of the microbiome, along with the role of metabolites in host-microbe interactions, necessitate the constant evolution of assay methods. To address this challenge, we propose a combined approach: a quantitative semi-targeted analysis (assay type 2 in Beger *et al*., 2024) of selected metabolites of interest, coupled with an untargeted metabolomics analysis. This combined approach makes it possible, firstly, to perform relative quantification of metabolites chosen with care and rationality in relation to the “fit-for-purpose” objective (see Figure 1a). Secondly, it enables us to carry out a fingerprint analysis, and above all to reanalyze the metabolites present in these matrices in the light of constantly evolving curated databases (see Figure 1b). Finally, the use of molecular networks makes it possible to generate new hypothesis concerning biomarker metabolites. These analyses need to encompass a wide physico-chemical range of metabolites while requiring minimum sample volume and resources, and downstream data processing workflows need to be as automated and rapid as possible. This is the approach we present in this article (see Figure 1).

**Figure 1.**
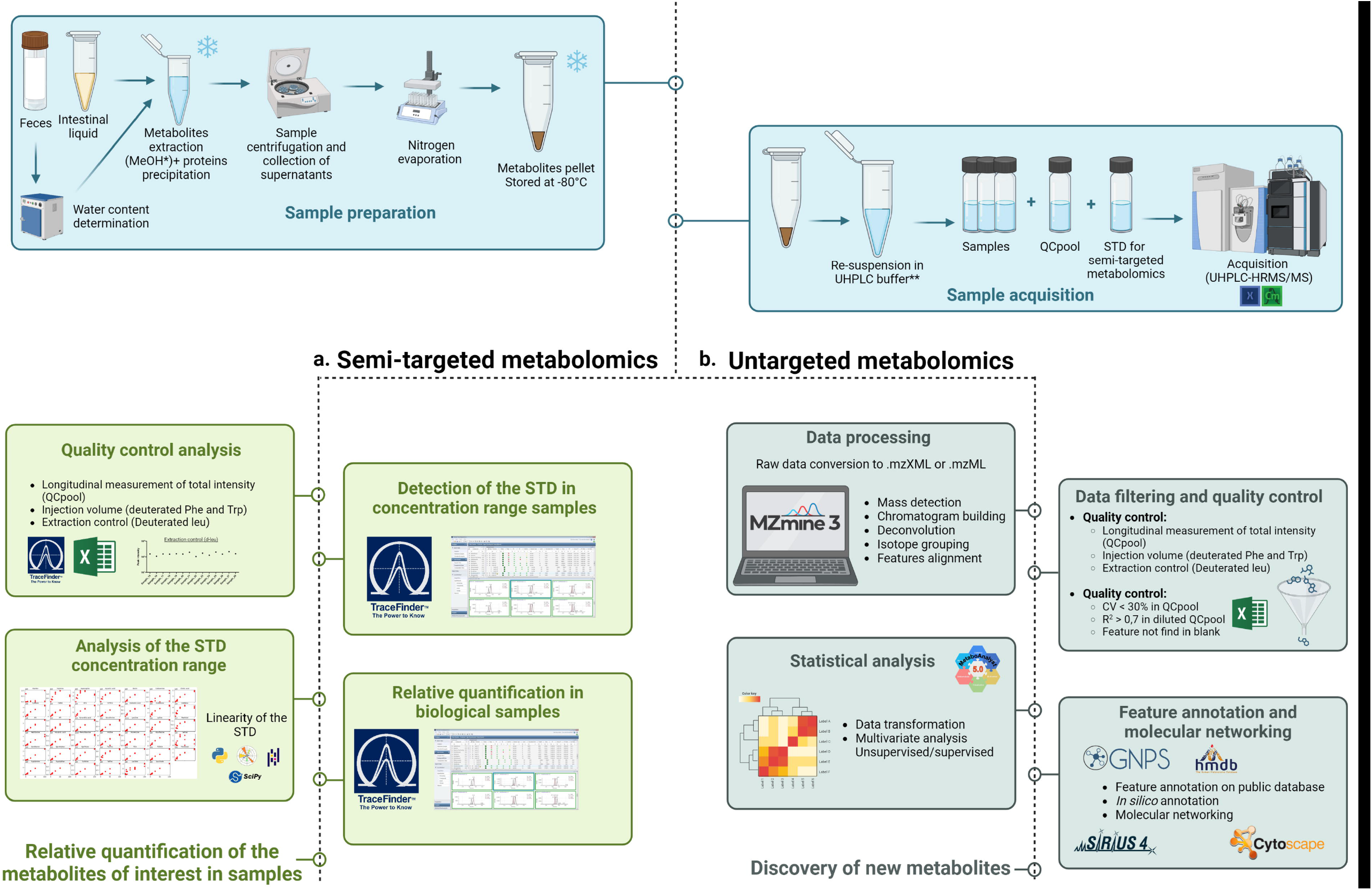
Workflow for the combined approach of semi-targeted and untargeted metabolomics to characterize small intestinal and fecal metabolomes. *Spike with d-leucine pike with d-tryptophan and d-phenylalanine.

### 3.2 Selection of the metabolites of interest for semi-targeted metabolomics approach

To select our metabolites of interest, we based our selection criteria on a workshop report by Mandal *et al*., which emphasized the importance of using human whole stool reference materials for metabolomics and metagenomics approaches [37]. The goal of the workshop was to identify metabolites for which evidence suggests relevance to health and disease, and to determine the appropriate steps to develop a reference material suited for this purpose. Given the rapidly evolving nature of gut microbiome science and the current state of knowledge, we searched the recent literature and added some metabolites based on the following criteria: 1) the metabolite impacts host health or is linked to a disease; 2) the metabolite is commonly found in feces; 3) the metabolite is part of a microbial metabolic pathway; and 4) the metabolite is commonly found in small intestinal samples. To be selected, a metabolite must meet at least two of these criteria. Mandal et al. proposed a list of microbially produced metabolites for the fecal microbiome, including SCFAs, BAs, amino acids, indole and its derivatives, and p-cresol [37]. We made our choices based on the criteria of the size of the metabolites (relevant for our instrument), chromatographic settings, and we select mainly the metabolites which are chemical-by-product of microbes (criteria number 3). A large proportion of the selected metabolites are amino acids, which are derived from food and serve as substrate for many microbial reactions [38], catabolism and biosynthesis of SCFAs [39] in the small intestine [39,40]. Free amino acids are relevant for health since they interact with cell receptors and modulate host functions [41]. SCFAs are the products of fiber degradation, and their concentrations increase in the distal part of the gut [17]. They are important microbial modulators of host functions [42–44]. Based on the recent literature, we added certain BAs. They were selected because they are produced by the host and can be metabolized by microbes. Primary conjugated BAs such as glycocholic acid (GCA) and taurocholic acid (TCA), were chosen as markers of the small intestine [8]. Secondary BAs, such as deoxycholic acid (DCA) and lithocholic acid (LCA), and tertiary BAs, such as ursodeoxycholic acid (UDCA), were included because they result from microbial transformation and are found in feces. BAs are major metabolites that contribute to health by interacting with various host cell receptors [45] and are associated with a wide range of functions, including immune [46,47] and metabolic (glucose and lipids homeostasis) processes [48]. Metabolites from tryptophan metabolism, such as indole, indole-3-propionic acid (IPA), indole-3-acetic acid (IAA), which are specifically synthetized by microbes, as well as kynurenic acid and kynurenine, were also added [49]. Other metabolites like polyamines (cadaverine, putrescine, spermine, spermidine) and also vitamins (biotin, riboflavin) were added due to their impact on host health and their production by gut microbes [23,50–52]. As a result, a list of 49 metabolites of great interest have been selected based on our criteria and are detailed in Supplementary data 1.

### 3.3 Development of the semi-targeted metabolomics strategy

The main objective was to develop a “fit-for-purpose” analytical assay, enabling us to accurately quantify previously selected metabolites in the small intestine and feces samples, and then validated the assay against the intended purpose [25]. We purchased and resuspended all the selected metabolites to create a collection of authentic chemical standards.

#### Determination of MS1 and MS2 spectra and RT

The first objective was to determine the MS1 and MS2 spectra for each metabolite and the RT depending on the chromatogram parameters. Each chromatogram and spectrum were manually analyzed using Xcalibur software (ThermoFisher Scientific). Table 1 shows the measured *m/z* ratio and RT for each metabolite. The RT ranged from 0.73 minute to 12.58 minutes. We obtained MS1 and MS2 spectra for all selected metabolites, except for indole, lithocholic acid, SCFAs, and p-cresol. We thus chose to remove these metabolites from our library. No MS2 spectra were found for lactic acid and alanine (fragments under 50 *m/z*). Additionally, we couldn’t distinguish CDCA, DCA and UDCA, all of which are dehydroxylated bile acids, due to their identical MS1 and close MS2 fragmentation.

**Table 1:** The measured *m/z ratio* and RT, ionization mode, M2 fragmentation, Coefficient of Variation (CV), correlation coefficient (R^2^), regression equation, linearity concentration ranges determined for each metabolite.

| Metabolite | Measured<br>*m/z*<br>(H<sup>+</sup> or H<sup>-</sup>) | RT | Ionization<br>mode | M2<br>fragmentation | CV | $R^2$ | Regression<br>equation | Concentration<br>range (μM) |
| --- | --- | --- | --- | --- | --- | --- | --- | --- |
| Alanine | 90.0557 | 1.23 | pos | no | 69.27 | 0.054 | $y = 2.525e+06 x + 9.708e+06$ | 1-100 |
| Arginine | 175.1192 | 1.17 | pos | yes | 11.01 | 0.927 | $y = 4.1e+07 x + 1.223e+08$ | 1-25 |
| Asparagine | 133.0611 | 1.21 | pos | yes | 12.65 | 0.867 | $y = 1.06e+07 x + 6.617e+07$ | 1-25 |
| Aspartic acid | 134.0449 | 1.17 | pos | yes | 13.6 | 0.853 | $y = 4.352e+06 x + 4.323e+07$ | 1-25 |
| Biotin | 245.0972 | 7.77 | pos | yes | 4.02 | 0.957 | $y = 2.88e+07 x + 1.41e+08$ | 1-50 |
| Cadaverine | 103.1237 | 0.79 | pos | yes | 69.7 | 0.66 | $y = 6.086e+06 x - 2.255e+07$ | 1-25 |
| Cholic acid | 409.2972 | 12.36 | pos | yes | 6.91 | 0.96 | $y = 2.525e+06 x + 9.708e+06$ | 1-50 |
| Cysteine | 122.0274 | 1.26 | pos | yes | 8.41 | 0.961 | $y = 6.136e+06 x + 2.812e+07$ | 1-50 |
| Gamma aminobutyric acid | 104.0711 | 0.97 | pos | yes | 6.27 | 0.962 | $y = 1.969e+07 x + 9.618e+07$ | 1-50 |
| Glutamic acid | 148.0614 | 0.96 | pos | yes | 7.35 | 0.958 | $y = 1.553e+07 x + 5.462e+07$ | 1-25 |
| Glutamine | 147.0765 | 1.23 | pos | yes | 5.55 | 0.926 | $y = 9.618e+06 x + 5.949e+07$ | 1-25 |
| Glycochenodeoxycholic acid | 450.3247 | 12.58 | pos | yes | 6.89 | 0.98 | $y = 1.849e+07 x + 1.079e+08$ | 1-100 |
| Glycocholic acid | 466.3191 | 11.34 | pos | yes | 6.56 | 0.943 | $y = 3.158e+07 x + 2.335e+08$ | 1-50 |
| Histidine | 156.0769 | 1.14 | pos | yes | 6.6 | 0.892 | $y = 1.267e+07 x + 8.603e+07$ | 1-25 |
| Indole-3-acetic acid | 176.0718 | 9.38 | pos | yes | 9.1 | 0.989 | $y = 7.027e+06 x + 2.277e+07$ | 1-100 |
| Indole-3-propionic acid | 190.0877 | 10.28 | pos | yes | 15.04 | 0.967 | $y = 1.209e+07 x + 4.067e+07$ | 1-100 |
| Kynurenic acid | 190.0511 | 7.08 | pos | yes | 5.84 | 0.986 | $y = 4.534e+07 x + 2.401e+08$ | 1-100 |
| Kynurenine | 209.0937 | 4.16 | pos | yes | 4.86 | 0.979 | $y = 5.537e+07 x + 2.816e+08$ | 1-100 |
| Lactic acid | 89.0231 | 1.44 | neg | no | 15.48 | 0.64 | $y = 4.354e+06 x + 9.303e+07$ | 1-25 |
| Leucine | 132.1029 | 1.86 | pos | yes | 5 | 0.986 | $y = 1.609e+08 x + 3.144e+08$ | 1-100 |
| Lysine | 147.1131 | 1.09 | pos | yes | 9.39 | 0.946 | $y = 1.401e+07 x + 4.41e+07$ | 1-100 |
| Mannitol | 183.0878 | 0.97 | pos | yes | 8.26 | 0.887 | $y = 1.892e+06 x + 1.464e+07$ | 1-25 |
| Methionine | 150.0596 | 1.51 | pos | yes | 15.78 | 0.975 | $y = 4.238e+07 x + 2.017e+08$ | 1-100 |
| Nicotinic acid | 124.0402 | 1.53 | pos | yes | 8.47 | 0.979 | $y = 5.53e+07 x + 3.164e+08$ | 1-100 |
| Phenylalanine | 166.0874 | 3.99 | pos | yes | 4.76 | 0.982 | $y = 9.785e+07 x + 4.205e+08$ | 1-100 |
| Proline | 116.0711 | 1.39 | pos | yes | 10 | 0.953 | $y = 5.868e+07 x + 8.56e+07$ | 1-50 |
| Putrescine | 89.108 | 0.79 | pos | yes | 6.76 | 0.988 | $y = 5.381e+06 x + 2.594e+07$ | 1-100 |
| Riboflavin | 377.148 | 7.69 | pos | yes | 6.26 | 0.97 | $y = 3.417e+07 x + 1.639e+08$ | 1-100 |
| Serine | 106.0504 | 1.15 | pos | yes | 52.87 | 0.798 | $y = 7.795e+06 x - 9.248e+06$ | 1-25 |
| Serotonin | 177.1036 | 4.45 | pos | yes | 3.52 | 0.995 | $y = 2.563e+07 x - 7.743e+07$ | 1-100 |
| Spermidine | 146.1663 | 0.73 | pos | yes | 6.11 | 0.989 | $y = 2.303e+07 x + 1.268e+08$ | 1-100 |
| Spermine | 203.2244 | 0.69 | pos | yes | 4.21 | 0.969 | $y = 4.583e+07 x + 1.808e+08$ | 1-100 |
| Succinic acid | 117.0182 | 1.81 | neg | yes | 7.04 | 0.992 | $y = 1.684e+07 x + 2.309e+07$ | 1-100 |
| Taurine | 126.0231 | 0.94 | pos | yes | 6.33 | 0.857 | $y = 2.319e+06 x + 2.589e+07$ | 1-25 |
| Taurochenodeoxycholic acid | 500.3078 | 11.33 | pos | yes | 6.29 | 0.985 | $y = 9.388e+06 x + 6.694e+07$ | 1-100 |
| Taurocholique acid | 516.3027 | 10.47 | pos | yes | 3.4 | 0.981 | $y = 2.35e+07 x + 8.876e+07$ | 1-100 |
| Threonine | 120.0658 | 1.22 | pos | yes | 10.25 | 0.933 | $y = 3.503e+06 x + 4.149e+07$ | 1-50 |
| Tryptamine | 161.1085 | 6.81 | pos | yes | 2.69 | 0.991 | $y = 7.745e+07 x + 3.109e+08$ | 1-100 |
| Tryptophan | 205.0986 | 6.44 | pos | yes | 2.69 | 0.987 | $y = 6.792e+07 x + 3.049e+08$ | 1-100 |
| Tyrosine | 182.0812 | 3.33 | pos | yes | 10.96 | 0.992 | $y = 5.853e+07 x + 1.236e+08$ | 1-100 |
| Valine | 118.0867 | 1.77 | pos | yes | 12.55 | 0.925 | $y = 5.845e+07 x - 7.895e+06$ | 1-100 |

#### Compound database creation

The compound database was built on TraceFinder4 software (ThermoFisher Scientific) from data collected previously. For this step, we built the compound database and associated a master method for quantification of the previously selected compound.

#### Repeatability of the metabolite’s detection

To ensure the reliability and accuracy of our semi-targeted metabolomics assay, we assessed repeatability, i.e. the ability to produce consistent results when repeated under identical conditions (same operator, same equipment, same reagents and samples, short time interval). For this, we measured the coefficient of variation (CV), the dispersion of measurements around the mean, which indicates the precision of the measurement. After injection of pooled metabolites 10 times in positive and negative ionization mode, the median CV of the detection was calculated. The peak intensity was obtained using TraceFinder4. The metabolites with a CV which were more than 30% were excluded. We found that 3 of the 41 selected metabolites had a CV > 30% (alanine, cadaverine and serine) (see Table 1). For medical biology analytical methods, an acceptable coefficient of variation threshold depends on the type of metabolite and the clinical application. In general, a CV of less than 15% is often considered acceptable for quantitative analysis. This is the case for the 39/41 metabolites tested (see table 1). The CV of each metabolite ranges from 2.69 to 15.78, with a median CV of 6.9% for the selected metabolites across replicates analyses.

#### Linearity of detection, determination of the linear dynamic concentration’s ranges and upper limits of quantification

The linearity of an assay method refers to its ability to produce results that are directly proportional to the concentration of the analyte within a specified range. In order to evaluate the linearity of the method, we prepared dilution series of pooled metabolites, knowing that the concentration of each metabolite varies from 1µM to 100µM. Then, the dilution series of pooled metabolites was injected three times, and the raw files were analyzed using TraceFinder4 to determine the peak area for each metabolite. After that, we plotted the calibration curve by graphing the instrumental responses against the concentrations. The linearity was evaluated using the linear correlation coefficient (R^2^) using Python (Scipy module), which should be close to 1 to indicate good linearity. Then, we determined the concentration range for each metabolite for which we got a R^2^>0.8 and manually placed the limit of linearity (see supplementary data 1). Finally, we calculated the R^2^ value for the obtained linear trend line (see Table 1). The regression equations could be used to evaluate the metabolite concentration in samples. If a plateau was observed after a certain concentration, it means that the upper limit of quantification (ULOQ) had been reached. Full data for linearity range are in Supplementary data 2. Over the 41 metabolites of interest tested, only 4 metabolites had an R^2^ < 0.8 (alanine, cadaverine, lactic acid, and serine), which is consistent with the repeatability results observed in the analysis of 3 of these compounds. The linear correlation coefficients (R^2^) of the selected compounds ranged from 0.853 to 0.995, 22/37 metabolites had a linear concentration ranging from 1 to 100 µM, 7/37 between 1 and 50 µM, and 8/37 between 1 to 25 µM.

After all analyses, 37 metabolites of interest had been validated using our combined approach. The whole process to set-up the semi-targeted metabolomics approach is described in Figure 2.

**Figure 2:**
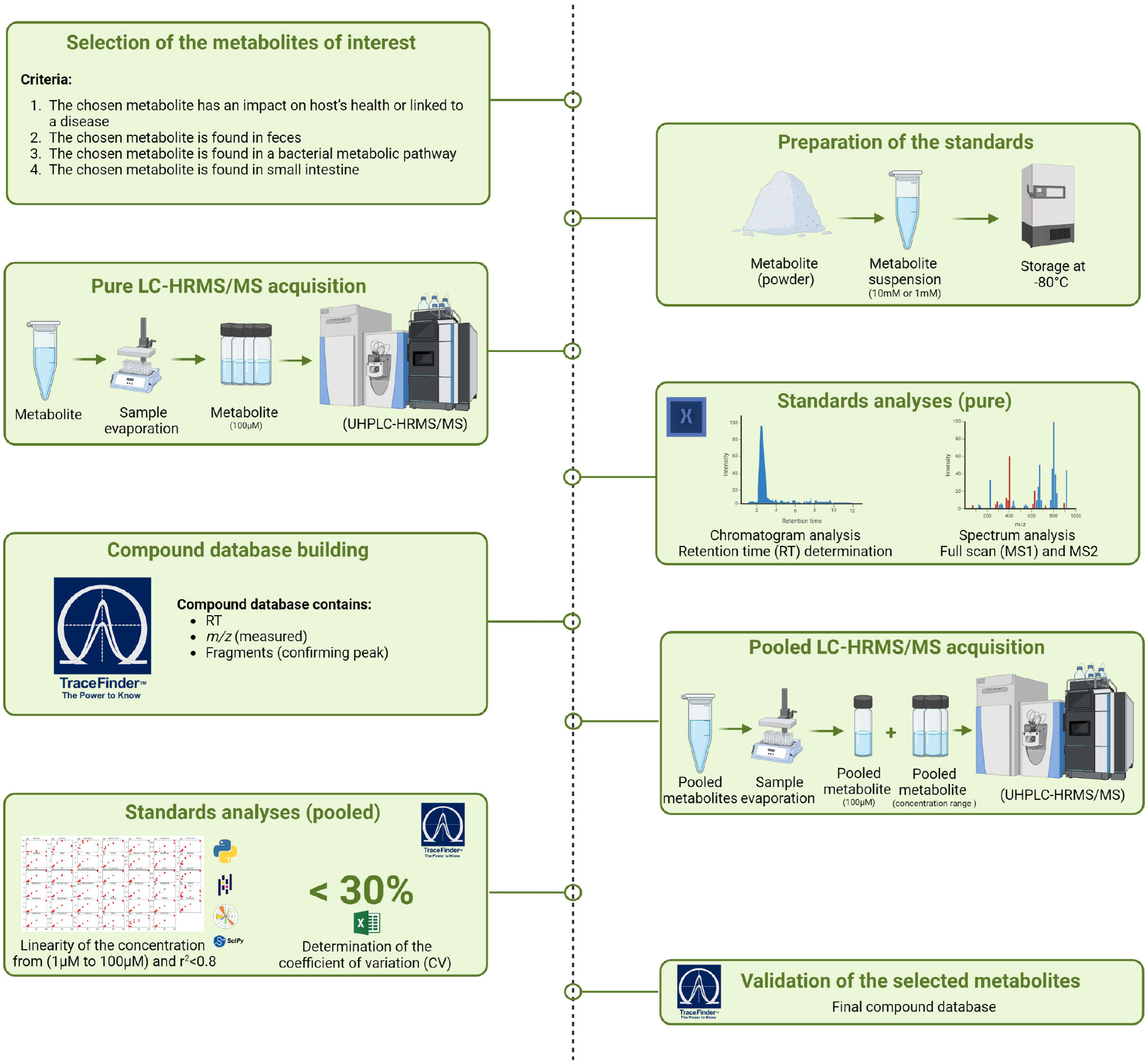
Development process of the semi-targeted metabolomics approach to characterize small intestinal and fecal metabolomes.

### 3.4 Development of the untargeted metabolomics strategy

The main objective of the untargeted approach is to determine the global metabolic profile of samples and allow the discovery of new metabolites. Data were processed following the pipeline described in Figure 1. After data processing on MZmine v3.0 and data filtering, features were statistically analyzed using MetaboAnalyst v6.0. The aligned feature lists were then exported for analysis using the Feature Base Molecular Networking tool [32] within the GNPS2 platform and features were annotated. GNPS annotations are based on MS1 and MS2 spectra. To increase the number of annotated features and to improve the consistency of the annotation, it’s also possible to perform in *silico* annotation. Based on MS2 spectra, fragmentation rules and probability, SIRIUS proposed annotations [33].

### 3.5 Proof of concept on clinical samples

In order to prove feasibility of our pipeline on biological samples, we tested our combined approach on samples from one participant of the clinical trial (NCT05477069) [14]. One small intestinal and fecal sample was extracted following the protocol described in materials and methods. For untargeted metabolomics, data were analyzed using MZmine3, Excel, GNPS2. The semi-targeted data were treated using TraceFinder4 and Python pipeline. The entire workflow for combined approach is described in Figure 1.

Thanks to the combined approach, we were able to observe distinct metabolomes between the small intestinal content and the fecal sample from the same donor (see Figure 3a). Indeed, 6155 features have been found in samples after MZmine processing, this includes all ion from such as multimers, different adducts and in/post source fragments. Following features filtration, 966 were specific to the small intestinal content, 2685 to the feces and 337 were shared across matrices. From the total filtrated features, 215 features were annotated with GNPS (<u>Link to GNPS job</u>). Thanks to the molecular networking, we identify BA new potential candidates in the samples (see Figure 3b). With the semi-targeted metabolomics, we consistently identified metabolites such as GCA, tryptophan, phenylalanine or leucine in samples. Using the chemical standard dilution series and linearity curves of these standards of interest (injected in parallel with the biological samples), we determined the concentrations of metabolites whose concentrations were within the linearity range. Among the 37 metabolites of interest, 28 were more abundant in the small intestinal content and 4 in the feces (see Figure 3c). Metabolites having an undetermined concentration (under the limit of detection) for one of the matrices are not represented on Figure 3c. For 9 metabolites (CA, GCA, glutamic acid, leucine, phenylalanine, proline, threonine, tyrosine and valine), the estimated concentrations were higher than the upper limit of quantification. Globally, amino acids and BAs were more abundant in the SI content. We evaluated that there was 1725 times more GCDCA in SI sample compared to the feces. These results are in accordance with previous studies [18,19]. Indeed, host derived BAs such as GCDCA are abundant in small intestinal before being transformed by microbes along the gut [13,47]. Metabolites that were more abundant in feces were biotin, nicotinic acid, aspartic acid, glutamic acid, kynurenine, serotonin, and tryptamine. Using molecular network, we demonstrate that we could find new bile acids in intestinal liquid sample compare to fecal samples. MS2 spectral matches to 68 bile acids were obtained by matching to the BILELIB19, a reference library available in GNPS dedicated to bile acids, increased to the annotations to 556. Matches to amino acid and other amine conjugated bile acids were also recovered which changed in distribution between small intestine and feces (Figure 3b).

**Figure 3:**
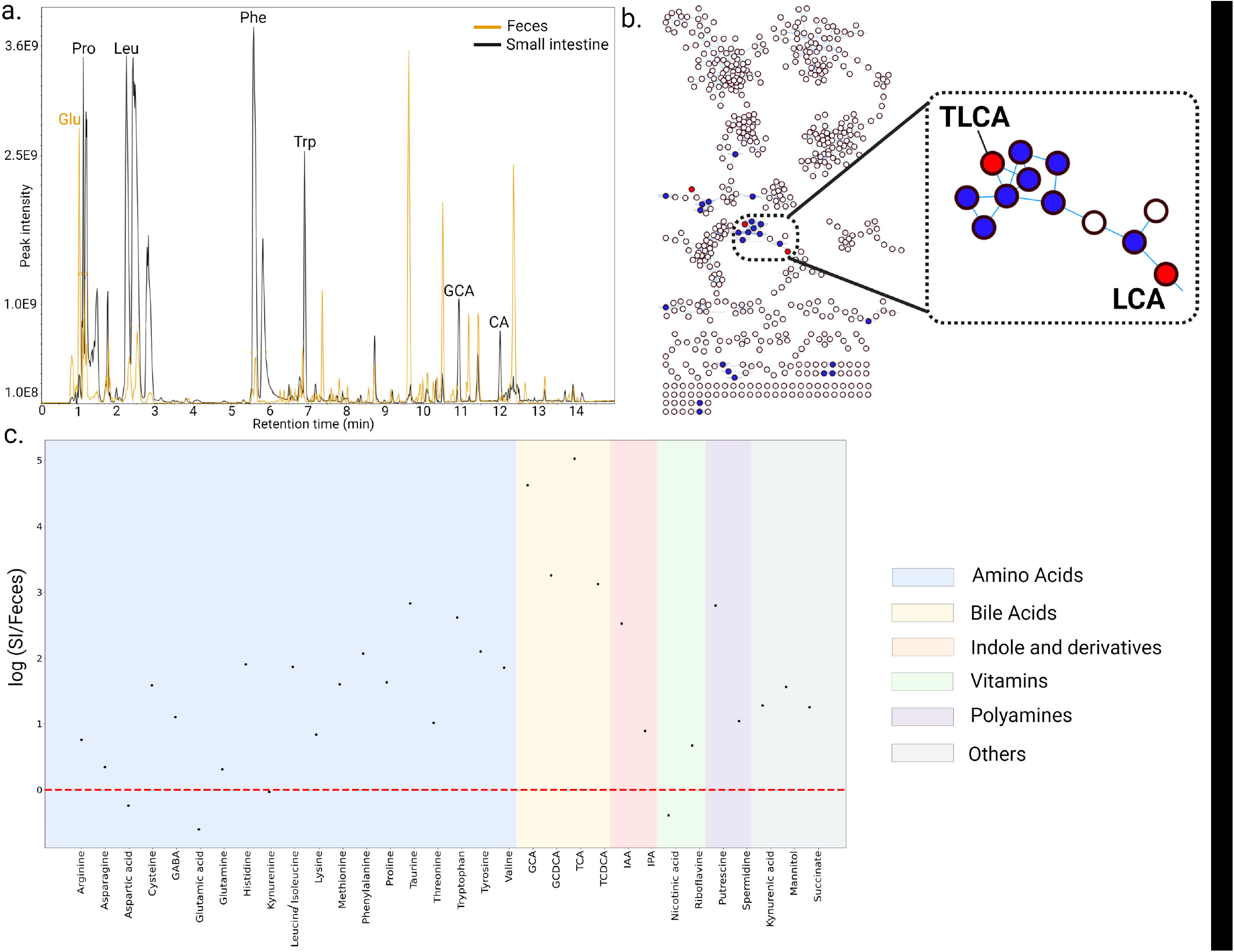
Proof of concept of the use of the combined metabolomics approach on small intestinal and fecal sample from one participant. (a) Global chromatograms from ll intestinal and fecal samples. (b) Molecular network of small intestinal and fecal samples, with a zoon on BA for the discovery of new molecules. (c) Ratios small stinal /feces sample for the 37 metabolites of interest in semi-targeted metabolomics. Metabolites are grouped depending on their classes (amino acids, BAs, vitamins, ole and derivatives, polyamines and others).

## 4 Conclusion

The combined approach presented in this article facilitates comprehensive metabolome profiling and the discovery of novel metabolites, including those derived from microbes. We determined the global metabolic profile of the samples and demonstrated the potential for discovering new molecules through molecular networking, employing an untargeted metabolomics strategy. Additionally, the semi-targeted approach enables high accurate determination of concentration of the selected metabolites and allows the relative quantification of metabolites across different samples. Moreover, we observed significant differences in metabolite ratio between small intestinal and fecal samples, with a higher abundance of metabolites in the small intestinal samples. We acknowledge that we have not validated all performance criteria of an analytical method aimed at determining the absolute concentration of metabolites. However, as stated at the beginning of the article, our objective was to develop a method that allows for the relative quantification of relevant metabolites, which we have carefully selected. This enables us to be robust in our measurements and to conclude on differences in concentrations between samples processed within the same batch. This workflow enhances gut microbiome research, providing deeper insights into the intricate relationships between gut microbiome and human health. Our method can be implemented with new metabolites. This would require analysis of the chemical standard of the new compound to add (i.e pure metabolite) to determine its RT, MS1 and MS2 spectra, followed by pooling of the latter to assess repeatability and linearity over a range of concentrations. Once validated, the metabolites can be added to the compound database.

Further studies, such as those by [53] and [54], have underscored the importance of integrating multiple metabolomics approaches to achieve a more holistic understanding of metabolic interactions and their implications for health and disease. By combining untargeted and semi-targeted methodologies, our approach aligns with these findings, offering a robust framework for advancing gut microbiome research and elucidating the complex interplay between microbial metabolites and host physiology.

## Author Contributions

AT and ALG conceived and designed research. AT, VC and CP conducted experiments. AT, MRM, CP, IM and ALG analyzed data. In particular, M.R.M designed all the python pipelines, the figures 3.c and S2. She helped for the TraceFinder treatment and data processing. AT and ALG wrote the manuscript. All authors read, reviewed and approved the manuscript.

## Ethical Statements

This research was approved by the CHUGA institutional review board and authorized after its filing with the CNIL according to the French procedure for a monocentric study and has been granted ethical approval by the Personal Protection Committee (23 February 2022 and 9 March 2023) and, by the French National Agency for the Safety of Medicines and Health Products (ANSM) (2 June 2022 and 20 March 2023), and it has formally been registered as a study (NCT05477069)[14].

## Data Availability

The metabolomics and metadata reported in this paper are available via MassIVE <u>MSV000095801</u>.

## Acknowledgments

We warmly thank Prof. D. Martin for the funding of a part of this research. We thank Dr. C. Trocmé, for her help on collecting samples during the clinical trial.

