## Supplementary data for "Untargeted and Semi-Targeted Metabolomics Approach for Profiling Small Intestinal and Fecal Metabolome Using High-Resolution Mass Spectrometry"

**Supplementary data 1: List of the candidate metabolites and the corresponding selection criteria**

| **Metabolite** | **Chemical class** | ***m/z*** | **InChi key** | **Selection criteria** | **Host effect** | **Literature** |
| --- | --- | --- | --- | --- | --- | --- |
| Alanine | Amino acid | 89,10 | QNAYBMKLOCPYGJ-REOHCLBHSA-N | 3 | Production of D-alanine by gut microbes | [1] |
| Arginine | Amino acid | 174,20 | ODKSFYDXXFIFQN-BYPYZUCNSA-N | 3 | Amino acid use as a substrate (biotransformation by microbial enzymes)  Arginine regulates the amino acids utilization by microbes derived from small intestine of pigs | [2]  [3] |
| Asparagine | Amino acid | 132,12 | DCXYFEDJOCDNAF-REOHCLBHSA-N | 3 | Amino acid use as a substrate (biotransformation by microbial enzymes) | [2] |
| Aspartic acid | Amino acid | 133,10 | CKLJMWTZIZZHCS-REOHCLBHSA-N | 3 | Precursor for SCFA (acetate) | [4] |
| Biotin | Vitamin | 244,3 | YBJHBAHKTGYVGT-ZKWXMUAHSA-N | 1 | Biotin is a coenzyme for five carboxylases, catalysing key steps in fatty acids, glucose and amino acids metabolisms.  Biotin deficiency is linked to intestinal inflammation and inflammatory bowel disease | [5]  [6] |
|  |  |  |  | 3 | 5% of the recommended daily allowance is provided by gut microbiota. The majority of *Bacteroidota*, *Fusobacteriota* and *Pseudomonadota* can synthesize biotin. | [7] |
| Butyric acid | SCFA | 88,11 | FERIUCNNQQJTOY-UHFFFAOYSA-N | 1 | SCFA have a panel of host cells receptors and influence host’s functions (immune, endocrine, barrier, …) | [8]  [9] |
|  |  |  |  | 2 | SCFA in feces (7.27 ± 4.42 μmol/g) | [10] |
|  |  |  |  | 3 | SCFA are produced in the gut by microbes (around 500-600mmol/day) | [8] |
| Cadaverine | Polyamine | 102,18 | VHRGRCVQAFMJIZ-UHFFFAOYSA-N 1 | 4 | Found in the small intestine lumen | [11] |
|  |  |  |  | 3 | Cadaverine is produced by bacteria by the decarboxylation of lysine | [12] |
|  |  |  |  | 1 | Cadaverine seems to reduce the proliferation and colony forming ability of breast cancer cells | [12] |
| Chenodeoxycholic acid | Bile acid | 392,57 | RUDATBOHQWOJDD-BSWAIDMHSA-N | 1 | Increased CDCA pool in IBD patient in stool  CDCA’s receptor in host’s cell like immune cells | [13]  [14]  [15] |
|  |  |  |  | 3 | BAs deconjugated by gut microbes | [16] |
|  |  |  |  | 4 | Conjugated primary BAs are secreted in the duodenum and deconjugated by gut microbes before reaching the colon | [16] |
| Cholic acid | Bile acid | 408,57 | BHQCQFFYRZLCQQ-OELDTZBJSA-N | 1 | Lower primary bile acids concentration in patient with constipation compared to healthy control and higher concentrations were detected in diarrhea patients. | [17] |
|  |  |  |  |  | Increased CA pool in IBD patient in stool | [13] |
|  |  |  |  | 4 | Conjugated primary BAs are secreted in the duodenum and deconjugated by gut microbes before reaching the colon | [16] |
|  |  |  |  | 3 | BAs are deconjugated by gut microbes (GCA and TCA to CA) | [18] |
| Cysteine | Amino acid | 121,16 | XUJNEKJLAYXESH-REOHCLBHSA-N | 3 | Amino acid use as a substrate (biotransformation by microbial enzymes) | [2] |
| Deoxycholic acid | Bile acid | 392,57 | KXGVEGMKQFWNSR-LLQZFEROSA-N | 1 | Increased CDCA pool in IBD patient in stool  CDCA’s receptor in host’s cell like immune cells | [13]  [15] |
|  |  |  |  | 2 | DCA is find in colon and feces | [19] |
|  |  |  |  | 3 | BAs deconjugated by gut microbes | [20] |
| Gamma aminobutyric acid | Anino acid | 103,12 | BTCSSZJGUNDROE-UHFFFAOYSA-N 1 | 3 | Bacteria producing GABA: genra *Bacteroides*, *Parabacteroides*, *Bifidobacterium* and *Escherichia* | [21] [22] |
|  |  |  |  | 1 | Major inhibitory neurotransmitter in Central nervous system 🡪 dysfunction in GABA metabolism involves anxiety and depression | [23] |
| Glutamic acid | Amino acid | 147,13 | WHUUTDBJXJRKMK-VKHMYHEASA-N | 1 | Glu important nutrient for the intestinal mucosal maintenance and is largely metabolized by enterocytes. Glu functions as a signaling molecule in the enteric nervous system and can modulates neuroendocrine reflexes. Possible link between Glu metabolism and Alzheimer disease and dementia. | [24]  [25] |
|  |  |  |  | 3 | Glu production by some lactic acid bacteria such as *Lactobacillus paracasei*  Amino acid use as a substrate (biotransformation by microbial enzymes) | [26]  [2] |
| Glutamine | Amino acid | 146,15 | ZDXPYRJPNDTMRX-VKHMYHEASA-N | 1 | Glutamine has an effect on gut integrity by promoting enterocyte proliferation, regulating tight junction proteins and conferring protection against apoptosis and cellular stress | [27] |
|  |  |  |  | 3 | Glutamine is a substrate for glutamate production by bacteria such as *Ruminococcus, Coprococcus* and *Dorea* | [28] |
| Glyco-chenodeoxycholic acid | Bile acid | 449,62 | GHCZAUBVMUEKKP-GYPHWSFCSA-N | 4 | Conjugated primary BAs produced by host and secreted in small intestine | [29]  [30] |
|  |  |  |  | 3 | GCDCA is metabolized by gut bacteria | [16] |
|  |  |  |  | 1 | Decrease of bile acid deconjugation in IBD patient | [14] |
| Glycocholic acid | Bile acid | 465,63 | RFDAIACWWDREDC-FRVQLJSFSA-N 1 | 4 | Conjugated primary BAs produced by host and secreted in small intestine | [29]  [30] |
|  |  |  |  | 3 | GCA is metabolized by gut bacteria | [16] |
|  |  |  |  | 1 | Decrease of bile acid deconjugation in IBD patient  Increase of GCA concentration in the bile of cholangiocarcinoma patients | [14]  [31] |
| Histidine | Amino acid | 155,16 | HNDVDQJCIGZPNO-YFKPBYRVSA-N | 3 | Amino acid use as a substrate (biotransformation by microbial enzymes) | [2]  [32] |
|  |  |  |  | 1 | Histidine is converted in histamine which is a proinflammatory factor | [33]  [34]  [35] |
| Indole-3-acetic acid | Indole and derivative | 175,18 | SEOVTRFCIGRIMH-UHFFFAOYSA-N 1 | 3 | Produced by gut bacteria from tryptophan | [36] |
|  |  |  |  | 1 | Aryl hydrocarbon receptor (AhR) ligand 🡪 modulation of the immune system development, function and maintenance. Influence gut permeability | [37]  [38] |
| Indole | Indole | 117,15 | SIKJAQJRHWYJAI-UHFFFAOYSA-N 1 | 1 | Indole has cell receptors (AhR) and module various host’s functions like endocrinal function. Indole stimulate L cells that produce glucagon-like peptide-1 (incretin). | [38]  [39] |
|  |  |  |  | 3 | Indole is produced from tryptophan metabolism by gut bacteria | [36]  [38] |
| Indole-3-propionic acid | Indole and derivative | 189,21 | GOLXRNDWAUTYKT-UHFFFAOYSA-N 1 | 1 | IPA can help to maintain the intestinal barrier by inducing cell proliferation  Neuroprotective activity | [40]  [41] |
|  |  |  |  | 3 | Produced by gut bacteria from tryptophan | [36] |
| Kynurenic acid | Aromatic acid | 189,17 | HCZHHEIFKROPDY-UHFFFAOYSA-N | 4 | Around 1,5 to 16µM in the lumen of rat’s small intestine | [42] |
|  |  |  |  | 3 | Some bacteria are able to produce kynurenic acid such as *Escherichia coli*. | [43] |
| Kynurenine | Amino acid | 208,21 | YGPSJZOEDVAXAB-QMMMGPOBSA-N | 1 | Increase concentration of kynurenine in colorectal cancer patient | [44] |
|  |  |  |  | 3 | Some bacteria can produce kynurenine such as *Pseudomonas fluorescens* | [45] |
| Lactic acid | Carboxylic acid | 90,08 | JVTAAEKCZFNVCJ-REOHCLBHSA-N 1 | 1 | Increase the neuronal activity by being an energy source for cells  Lactate likely linked to GABA and boost mice memory  Stimulate intestinal stem-cells proliferation | [46]  [47]  [48] |
|  |  |  |  | 3 | Produced by bacteria from carbohydrates | [49] |
|  |  |  |  | 4 | Lactate is found in small intestinal content | [50]  [49] |
| Leucine | Amino acid | 131,17 | ROHFNLRQFUQHCH-YFKPBYRVSA-N | 1 | Promote intestinal health by increasing the secretion of IgAs | [51] |
|  |  |  |  | 3 | Amino acid use as a substrate (biotransformation by microbial enzymes) | [2] |
| Lithocholic acid | Bile acid | 376,57 | SMEROWZSTRWXGI-HVATVPOCSA-N | 1 | LCA binds to cell receptors and induces cytokines production and increases differentiation of regulatory T cells (reduce inflammation). Downregulation of interleukin 1 by LCA leads to vasodilatation in endothelial cells.  LCA may be consider as carcinogen in high concentration by inducing oxidative stress and DNA damages. | [52]  [53] |
|  |  |  |  | 2 | LCA is found in feces | [54] |
|  |  |  |  | 3 | LCA is a secondary BA produced by gut microbes such as *Clostridium* and *Eubacterium* | [53] |
| Lysine | Amino acid | 146,19 | KDXKERNSBIXSRK-YFKPBYRVSA-N | 3 | Amino acid use as a substrate (biotransformation by microbial enzymes)  Precursor for SCFA (acetate and butyrate)  Lysine can be produced by gut bacterial | [2]  [4] |
|  |  |  |  | 4 | Lysine is produced by small intestinal microbes | [55] |
| Mannitol | Polyol | 182,17 | FBPFZTCFMRRESA-KVTDHHQDSA-N 1 | 1 | Associate with Alzheimer disease and cognitive decline (increased in patients) in plasma | [56] |
|  |  |  |  | 3 | Mannitol is a prebiotic and a substrate for gut microbes  Production of mannitol from certain lactic acid bacteria | [57]  [58] |
| Methionine | Amino acid | 149,21 | FFEARJCKVFRZRR-BYPYZUCNSA-N | 1 | The methionine metabolism is implicated in inflammation, cancer and neurological diseases. Decrease of methionine serum and fecal levels in fatigue patient with quiescent IBD | [59]  [60] |
|  |  |  |  | 3 | Amino acid use as a substrate (biotransformation by microbial enzymes)  First substrate for methionine metabolism | [2]  [60] |
| Nicotinic acid | Vitamin | 123,10 | PVNIIMVLHYAWGP-UHFFFAOYSA-N | 3 | 27% of the recommended daily allowance is provided by gut microbiota. Can be synthesized by *Bacillota* and *Actinomycetota* and some *Pseudomonadota* | [7] |
| P-cresol | Phenol | 108.13 | IWDCLRJOBJJRNH-UHFFFAOYSA-N | 1 | P-cresol is correlated with chronic kidney disease  In mice, it promotes small intestinal transit | [61]  [62] |
|  |  |  |  | 3 | P-cresol is a microbially derived metabolite from the tyrosine metabolism | [62] |
| Phenylalanine | Amino acid | 165,19 | COLNVLDHVKWLRT-QMMMGPOBSA-N | 3 | Phenylalanine (Phe) can be produced by gut microbes and is the basis of Phe metabolism | [33] |
| Proline | Amino acid | 115,13 | ONIBWKKTOPOVIA-BYPYZUCNSA-N | 3 | Amino acid use as a substrate (biotransformation by microbial enzymes) | [2] |
| Putrescine | Polyamine | 88,15 | KIDHWZJUCRJVML-UHFFFAOYSA-N 1 | 4 | Putrescine is the most abundant polyamine in the small intestine | [63] |
|  |  |  |  | 2 | Putrescine in stool in healthy group around 800µM | [64] |
|  |  |  |  | 3 | Putrescine can be produced by gut microbes | [65] |
| Riboflavin | Vitamin | 376,36 | AUNGANRZJHBGPY-SCRDCRAPSA-N | 1 | Plays a key role in the energy and redox metabolism in both microbes and host  Riboflavin has an anti-inflammatory and antioxidant effects | [66] |
|  |  |  |  | 3 | Bacteria from *Bacteroidota* and *Fusobacteriota* and some from *Pseudomonadota* and *Bacillota* phyla contain all essential functions to synthesized riboflavin | [7] |
| Serine | Amino acid | 105,09 | MTCFGRXMJLQNBG-REOHCLBHSA-N | 3 | Amino acid use as a substrate (biotransformation by microbial enzymes) | [2] |
| Serotonin | Indole and derivative | 176,21 | QZAYGJVTTNCVMB-UHFFFAOYSA-N 1 | 1 | Modulate the intestinal motility and secretion  Hyperserotonemia in 25% of children with autism spectrum disorders  Serotonin level is decreased in metabolic syndrome and is negatively correlated with body mass index and body fat | [46]  [67]  [68] |
|  |  |  |  | 3 | Serotine can be produced by gut bacteria like *Klebsiella, Hafnia, Lactococcus.* | [69] |
| Spermidine | Polyamine | 145,25 | ATHGHQPFGPMSJY-UHFFFAOYSA-N | 1 | Effects on cardio-protection, tumor suppression, immune modulation, neuroprotection. Improve gut barrier integrity | [70]  [71] |
|  |  |  |  | 3 | Produce by gut microbes | [70] |
| Spermine | Polyamine | 202,34 | PFNFFQXMRSDOHW-UHFFFAOYSA-N | 1 | Anti-inflammatory effects by the inhibition of LPS induced expression of proinflammatory cytokines by monocytes and macrophages | [72] |
|  |  |  |  | 3 | Bacteria can produce spermine from putrescine | [63] |
| Succinic acid | Dicarboxylic acid | 118,09 | KDYFGRWQOYBRFD-UHFFFAOYSA-N 1 | 1 | Associate with Alzheimer disease and cognitive decline (increased in patients) | [56] |
|  |  |  |  | 3 | Product by *Parabacteroides distasonis* can reduce hyperglycemia and limit obesity | [73] |
| Taurine | Amino acid | 125,15 | XOAAWQZATWQOTB-UHFFFAOYSA-N | 1 | Taurine metabolism downregulated in Parkinson’s disease mice model. Taurine supplement has a protective effect | [74] |
|  |  |  |  | 3 | Taurine is a source of energy, source of sulphur (form food and conjugated bile acids) | [75] |
| Tauro-chenodeoxycholic acid | Bile acid | 499,71 | BHTRKEVKTKCXOH-BJLOMENOSA-N | 4 | Conjugated primary BAs produced by host and secreted in small intestine | [29]  [30] |
|  |  |  |  | 1 | Decrease of bile acid deconjugation in IBD patient  Decrease of TCDCA concentration in the bile of cholangiocarcinoma patients | [14]  [31] |
|  |  |  |  | 3 | TCDCA is metabolized by gut microbes | [16] |
| Taurocholic acid | Bile acid | 515,70 | WBWWGRHZICKQGZ-HZAMXZRMSA-N | 4 | Conjugated primary BAs produced by host and secreted in small intestine  Higher level of TCA in small intestine compared to stool | [29]  [30] |
|  |  |  |  | 1 | Decrease of bile acid deconjugation in IBD patient | [14] |
|  |  |  |  | 3 | TCA is metabolized by gut microbes | [16] |
| Threonine | Amino acid | 119,12 | AYFVYJQAPQTCCC-GBXIJSLDSA-N | 3 | Amino acid use as a substrate (biotransformation by microbial enzymes)  Precursor for SCFA (acetate, butyrate and propionate) | [2]  [4] |
|  |  |  |  | 4 | Microbial production by small intestinal microbes | [55] |
| Tryptamine | Indole and derivative | 160,21 | APJYDQYYACXCRM-UHFFFAOYSA-N 1 | 3 | Some bacteria such as *Ruminococcus, Blautia, Lactobacillus* and *Clostridium* are able to convert tryptophan to tryptamine | [76] |
| Tryptophan | Amino acid | 204,23 | QIVBCDIJIAJPQS-VIFPVBQESA-N | 1 | Many diseases linked to tryptophan metabolism disruption (e.g. IBD; IBS, metabolic syndrome) | [38] |
|  |  |  |  | 3 | Tryptophan is the precursor of kynurenine, serotonin and indole pathways  Amino acid use as a substrate (biotransformation by microbial enzymes) | [36]  [2] |
| Tyrosine | Amino acid | 181,19 | OUYCCCASQSFEME-QMMMGPOBSA-N | 3 | Amino acid use as a substrate (biotransformation by microbial enzymes) | [2]  [62] |
| Ursodeoxycholic acid | Bile acid | 392,57 | RUDATBOHQWOJDD-UZVSRGJWSA-N | 1 | UDCA found in higher level in IBS feces compared to healthy subjects | [14] |
|  |  |  |  | 2 | UDCA is found in feces | [14] |
|  |  |  |  | 3 | UDCA is a tertiary BA produced by microbes | [77] |
| Valeric acid | SCFA | 102,13 | NQPDZGIKBAWPEJ-UHFFFAOYSA-N 1 | 1 | Valeric acid is a histone deacetylase (HD) inhibitor. The overexpression of HD linked to various diseases | [78] |
|  |  |  |  | 3 | Valeric acid derives from microbial metabolism | [79] |
| Valine | Amino acid | 117,15 | KZSNJWFQEVHDMF-BYPYZUCNSA-N | 3 | Amino acid use as a substrate (biotransformation by microbial enzymes) | [2] |

Criteria:

(1): the metabolite impacts host health or is linked to a disease

(2): the metabolite is commonly found in feces

(3): the metabolite is part of a microbial metabolic pathway

(4): the metabolite is commonly found in small intestinal samples

**Supplementary data 2: concentration range and the R^2^ determined for each metabolite**
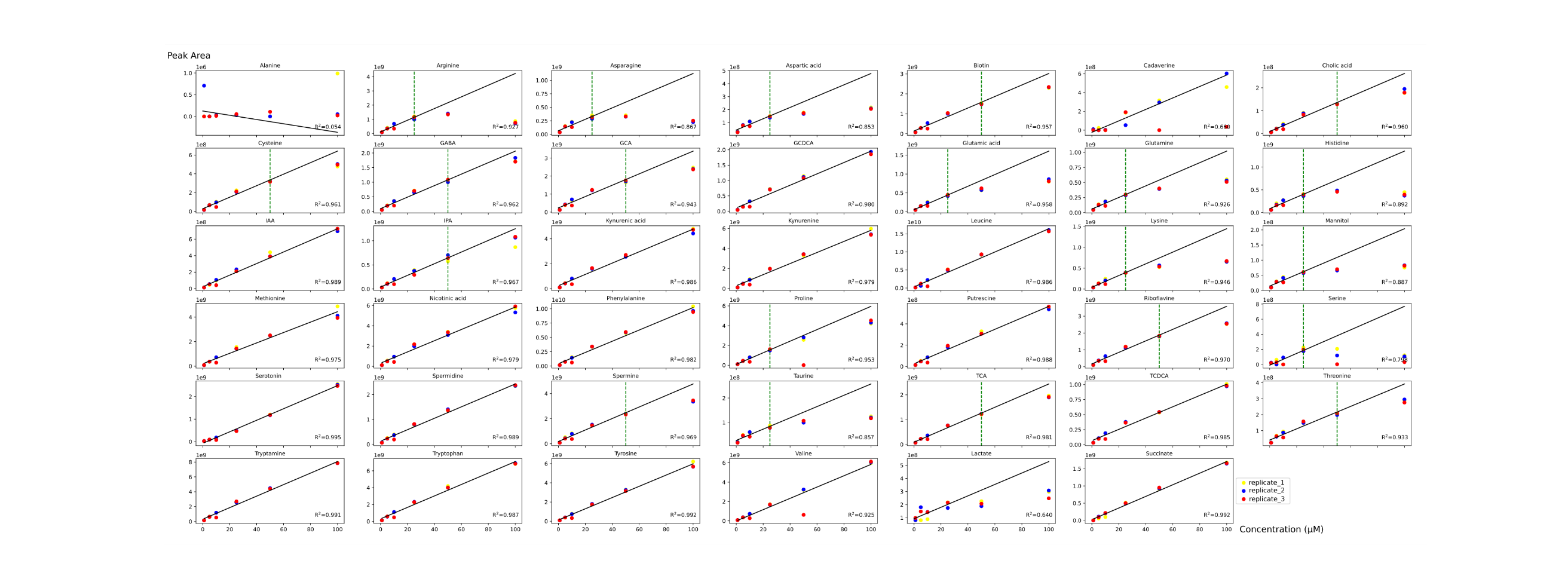

GABA: gamma aminobutyric acid, GCA: glycocholic acid, GCDCA: glycochenodeoxycholic acid, IAA: indole-3-acetic acid, IPA: indole-3-propionic acid, TCA: taurocholic acid, TCDCA: taurochenodeoxycholic acid. The green line corresponds to the end of the linearity in the tested concentration range. When there is not a green line, the validated concentration range is 1-100µM. The different point colors correspond to the triplicates of injection.
